# Huntington’s disease age at motor onset is modified by the tandem hexamer repeat in *TCERG1*

**DOI:** 10.1101/2021.07.16.452643

**Authors:** Sergey V. Lobanov, Branduff McAllister, Mia McDade-Kumar, G. Bernhard Landwehrmeyer, Michael Orth, Anne E. Rosser, for the REGISTRY Investigators of the European Huntington’s disease network, Jane S. Paulsen, for the Predict-HD study, Jong-Min Lee, Marcy E. MacDonald, James F. Gusella, Jeffrey D. Long, Mina Ryten, Nigel Williams, Peter Holmans, Thomas H. Massey, Lesley Jones

**Affiliations:** Medical Research Council Centre for Neuropsychiatric Genetics and Genomics, Cardiff University, United Kingdom; Department of Neurology, University of Ulm, Ulm, Germany; Department of Old Age Psychiatry and Psychotherapy, Bern University, Bern, Switzerland; Swiss Huntington’s Disease Centre, Siloah, Gümligen, Switzerland; School of Biosciences, Cardiff University, Cardiff, CF10 3AX, United Kingdom; University of Iowa, Iowa City, Iowa, United States, 52242; Dept of Neurology, University of Wisconsin, Madison, WI53705, USA; Molecular Neurogenetics Unit, Center for Genomic Medicine, Massachusetts General Hospital, Boston MA 02114, USA; Department of Neurology, Harvard Medical School, Boston MA 02115, USA; Medical and Population Genetics Program, the Broad Institute of M.I.T. and Harvard, Cambridge MA 02142, USA; Department of Genetics, Blavatnik Institute, Harvard Medical School, Boston MA 02115, USA; Great Ormond Street Institute of Child Health, Genetics and Genomic Medicine, University College London, London, United Kingdom; NIHR Great Ormond Street Hospital Biomedical Research Centre, University College London, London, United Kingdom; UK Dementia Research Institute at Cardiff, Cardiff University, United Kingdom

**Keywords:** Huntington’s disease, *TCERG1*, age at onset, short tandem repeat, quasi-tandem repeat, single nucleotide variant, whole exome sequencing

## Abstract

**Background:** Huntington’s disease is caused by an expanded CAG tract in *HTT*. The length of the CAG tract accounts for over half the variance in age at onset of disease, and is influenced by other genetic factors, mostly implicating the DNA maintenance machinery. We examined a single nucleotide variant, rs79727797, on chromosome 5 in the *TCERG1* gene, previously reported to be associated with Huntington’s disease and a quasi-tandem repeat (QTR) hexamer in exon 4 of *TCERG1* with a central pure repeat.

**Methods:** We developed a novel method for calling perfect and imperfect repeats from exome sequencing data, and tested association between the QTR in *TCERG1* and residual age at motor onset (after correcting for the effects of CAG length in the *HTT* gene) in 610 individuals with Huntington’s disease via regression analysis.

**Results:** We found a significant association between age at onset and the sum of the repeat lengths from both alleles of the QTR (p = 2.1×10^−9^), with each added repeat hexamer reducing age at onset by one year (95% confidence interval [0.7, 1.4]). This association explained that previously observed with rs79727797.

**Conclusions:** The association with age at onset in the genome-wide association study is due to a QTR hexamer in *TCERG1*, translated to a glutamine/alanine tract in the protein. We could not distinguish whether this was due to cis-effects of the hexamer repeat on gene expression or of the encoded glutamine/alanine tract in the protein. These results motivate further study of the mechanisms by which *TCERG1* modifies onset of HD.

## Background

Huntington’s disease (HD) is an autosomal dominant neurodegenerative disorder caused by an expanded CAG tract in exon 1 of the huntingtin gene (*HTT*). It typically manifests as a progressive movement disorder, often associated with debilitating cognitive, psychiatric and behavioural problems [1]. Symptoms usually start in mid-life, progressing over 10-30 years to dementia and premature death [2]. The CAG tract is polymorphic in the normal population with 6-35 CAGs, with 36 or more CAGs in HD subjects. There is an inverse correlation between CAG tract length and age at onset of disease symptoms, accounting for up to 70% of the variance in age at onset [3–6]. Genome-wide association studies (GWAS) have shown that other genetic variants also influence age at onset of HD, including variants in genes in DNA damage repair pathways and sequence variants in the CAG tract [7–9]. The most recent genetic modifier GWAS in HD (GeM-GWAS) [8] revealed 21 independent signals at 14 loci. We observed that one of the significant loci on chromosome 5 (5BM1) contained *TCERG1*, the only putative genetic modifier of HD onset in the GWAS to have been previously reported [10,11]. The 5BM1 locus (146 Mbp; hg19) has one significant single nucleotide variant (SNV), rs79727797 (p = 3.8 × 10^−10^), with each minor allele conferring 2.3 years later onset of HD than expected from the subjects’ CAG repeat length. SNV rs79727797 is within the *TCERG1* gene and very close to the tandem repeat locus (Fig. 1A) previously implicated in modifying HD age at onset [10,11].

**Fig. 1.**
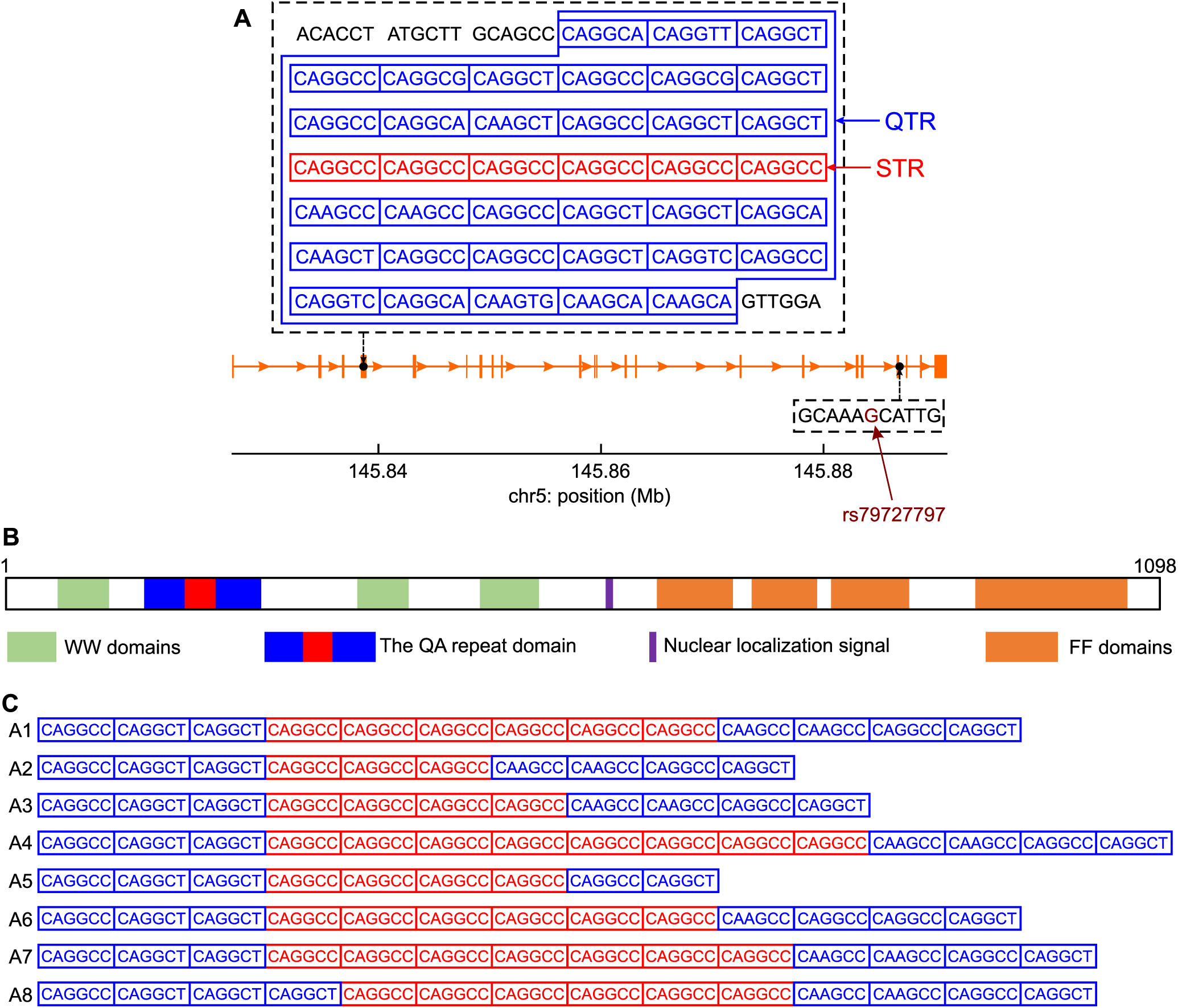
The relationship of rs79727797 to the CAGGCC hexanucleotide short tandem repeat in *TCERG1*. **a** The sequence of the tandem repeat region in exon 4 of *TCERG1* (orange). The blue polygon bounds quasi-tandem repeat (QTR) the central part of which contains pure repeat, CAGGCC hexanucleotide short tandem repeat (STR). **b** The *TCERG1* protein domains and location of the repeat tract. **c** The variant alleles seen at the tandem repeat locus arranged in descending order of prevalence.

*TCERG1* (Transcriptional Elongation Regulator 1; previously known as CA150) protein couples transcriptional elongation and splicing, regulating the expression of many genes [12,13]. It is highly conserved across human and mouse (97.8% identity between proteins). In humans, TCERG1 is extremely intolerant to loss of function variants (observed/expected variants = 0.13, 90% CI 0.07 – 0.23) and is in the 5% of genes most intolerant of amino acid missense substitutions, (observed/expected variants = 0.61, 90% CI 0.56 – 0.67) [14]. *TCERG1* binds to HTT and its expression can rescue mutant HTT neurotoxicity in rat and mouse model systems [15]. *TCERG1* contains a repeat tract of 38 tandem hexanucleotides: a central perfect short tandem repeat (STR) of (CAGGCC)_6_ embedded in a larger imperfect hexanucleotide ‘quasi’ tandem repeat (QTR; Fig. 1A,B; chr5:145,838,546-145,838,773 on hg19). The whole tract is translated in *TCERG1* protein as an imperfect 38 glutamine/alanine (QA) repeat interrupted with occasional valines (V; Additional file 1: Fig. S1).

Previously, a study of 432 American HD patients showed a nominally significant association of earlier onset with longer QTR length in *TCERG1* (p = 0.032, not corrected for multiple testing) [10]. A study of 427 individuals from Venezuelan HD kindreds [11] testing 12 polymorphisms previously associated with HD gave a p-value of 0.07 (not corrected for multiple testing) comparing the 306bp allele (corresponding to the reference 38-repeat QTR) with all other alleles for association with age at onset. Neither study tested the effects of repeat length directly, instead inferring it from the length of the amplified PCR products, including the flanking primer sequences.

We directly determined the repeat tract sequence in *TCERG1* in 610 HD patients by using short-read exome-sequencing data [1]. We then assessed the association of repeat alleles with age at onset of HD. We used a subset of 468 individuals for whom SNV data were available to test whether the rs79727797 variant was tagging the tandem repeat in *TCERG1* and whether the tandem repeat was likely to be the functional variant involved in modifying HD age at onset.

## Results

### Alleles observed at the *TCERG1* hexamer repeat

Subjects came from the REGISTRY [16] and Predict-HD [17] studies, and in Registry were individuals with the largest difference between their observed age at motor onset and that expected given their CAG repeat length, and in PREDICT those with the most extreme phenotype given their CAG repeat length, as in McAllister et al. [1].

The 38-unit QTR locus is in exon 4 of *TCERG1* and SNV rs79727797 just 3’ to exon 19, separated by 50 kbp (Fig. 1A). The length of the QTR is polymorphic and we identified eight different alleles, mostly varying by central STR length (Fig. 1C). The reference allele (A1), with a central (CAGGCC)_6_ STR, is by far the most common allele, representing 91.3% of all alleles sequenced in our study (Table 1). Alternative alleles with central STRs of different lengths were observed (Fig. 1C), of which the most common was (CAGGCC)_3_ (4.1% of alleles; A2, Table 1). This three-repeat allele is in linkage disequilibrium with the minor allele of rs79727797: in our cohort, correlation between the SNV and allele A2 is 99%.

**Table 1.** Hexanucleotide repeat allele frequencies in *TCERG1* (QTR = quasi-tandem repeat; STR = short tandem repeat)

| Allele | QTR length |  | STR length |  | Number of alleles | Allele frequency (%) |
| --- | --- | --- | --- | --- | --- | --- |
| | N | $\Delta N$ | N | $\Delta N$ | | |
| A1 | 38 | 0 | 6 | 0 | 1114 | 91.31 |
| A2 | 35 | -3 | 3 | -3 | 50 | 4.10 |
| A3 | 36 | -2 | 4 | -2 | 28 | 2.30 |
| A4 | 40 | +2 | 8 | +2 | 24 | 1.97 |
| A5 | 34 | -4 | 4 | -2 | 1 | 0.08 |
| A6* | 38 | 0 | 6 | 0 | 1 | 0.08 |
| A7 | 39 | +1 | 7 | +1 | 1 | 0.08 |
| A8 | 39 | +1 | 6 | 0 | 1 | 0.08 |

\* Allele A6 differs from the reference one, A1, by a synonymous SNV (see Fig. 1B).

### Association with age-at onset of HD

The distribution of genotypes observed in our study is given in Fig. 2. We tested for association between residual age at onset of HD and the QTR length. As there are two alleles, we examined the association with residual age at onset of the larger or smaller repeat length, the sum of repeat lengths, and the difference between repeat lengths in each patient. We consistently found higher levels of significance in the association between residual age at onset and the sum of the repeat lengths than in the associations with the difference between repeat lengths, or maximum or minimum repeat lengths in each individual (Additional file 1: Table S1). The association of the sum of the QTR lengths from both alleles with residual age at onset was genome-wide significant (p = 5.0×10^−9^ and 2.0×10^−8^ without and with multiple testing correction, respectively) (Additional file 1: Table S1). Logistic regression analyses using the extremes of the residual age at onset showed a similar pattern. The relationship between the sum of the hexamer repeats and the residual age at onset in HD is illustrated in Fig. 3 (see also Additional file 1: Fig. S2 for equivalent analyses of STR). Panels A-C show that subjects with extreme late onset have more copies of the shorter alleles than those with extreme early onset, and this difference becomes more pronounced as the extremes become greater. The negative correlation between the sum of QTR lengths in an individual and residual age at onset of HD is shown in Fig. 3D, with one year earlier HD onset for each added repeat hexamer (black dashed line in Fig. 3D, 95% confidence interval [0.7, 1.4]). We estimated the QTR effect size using the regression with selection analysis described in Additional file 1: Supplementary Methods. Since our HD cohort mainly contains age at onset extremes, the linear regression analysis (grey dashed line in Fig. 3D) overestimates the QTR effect size, giving 2.75 years earlier for each added hexamer. However, it can be used for comparison of the association significance between different models because it provides approximately the same *p*-value as the regression with selection analysis (Additional file 1: Table S2). Additional file 1: Table S2 shows a significant negative association between age at onset and the sum of QTR repeat lengths in both the REGISTRY and Predict-HD samples. Notably, (Table S2), the effect size estimated in the REGISTRY sample using regression with selection (0.98 years earlier onset for each added hexamer) is similar to that observed in the Predict-HD sample, where the selection is less extreme (1.26 years earlier onset for each added hexamer). This is an indication that applying regression with selection has successfully corrected for the bias in effect size induced by the extreme onset selection in the REGISTRY sample. The associations are slightly less significant when the pure hexamer repeat length is used rather than the full repeat: p = 6.5 × 10^−9^ for linear regression (Additional file 1: Table S3). However, the sample size is relatively small, and a larger sample would be needed to establish whether there is any significant difference between these results.

**Fig. 2.**
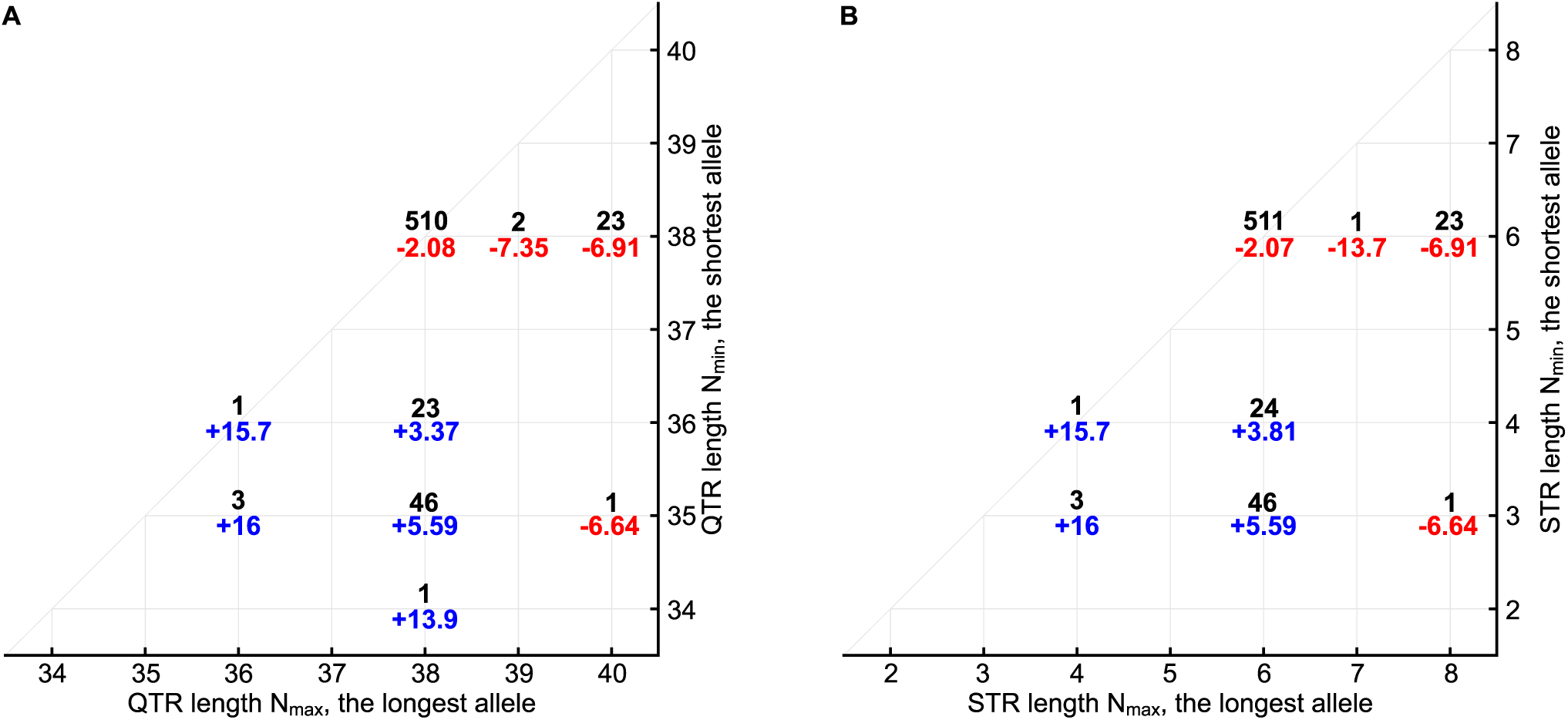
*TCERG1* tandem repeat genotype counts and associated mean residual ages at onset. **a** Quasi-tandem repeat (QTR) genotypes; **b** Short tandem repeat (STR) genotypes. Black numbers mark genotype counts. Red and blue numbers indicate mean residual ages at onset for individual genotypes, early onset in red, late onset in blue.

**Fig. 3.**
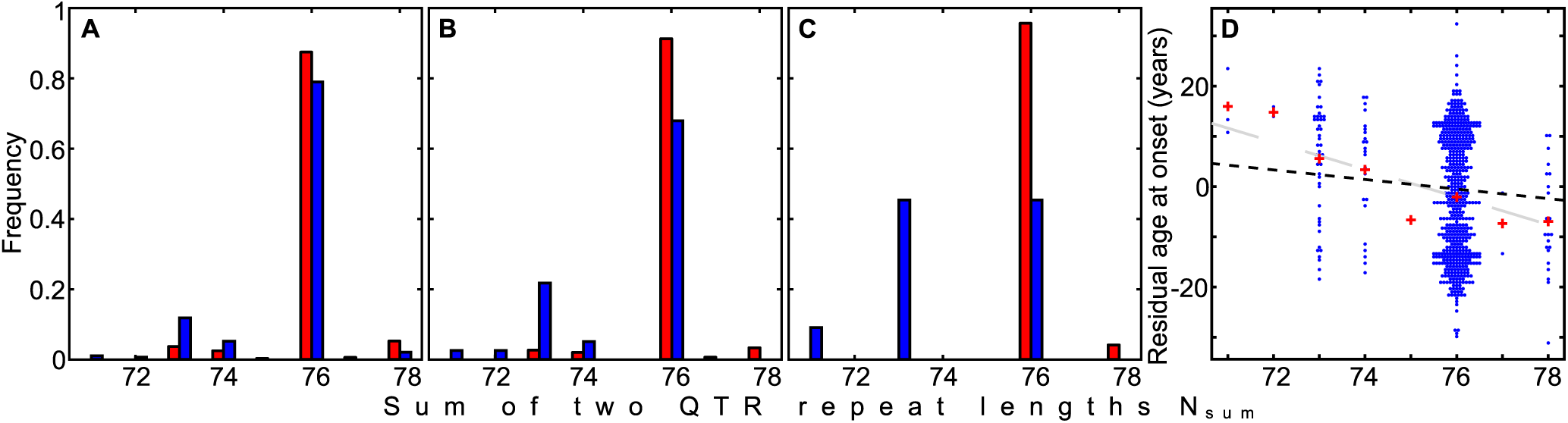
The relationship between hexanucleotide short tandem repeat (STR) length and residual age at onset of HD. **a-c** Histograms showing distribution of the sum of two STR repeat lengths N_sum_ = N_min_ + N_max_ for the groups with early (red, R<-R_thr_) and late (blue, R>R_thr_) onsets. The panels **a, b**, and **c** correspond to the residual age at onset threshold R_thr_ of 0, 13, and 20 years, respectively. **d** Association of the sum of two STR repeat lengths N_sum_ with the residual age at onset for the entire HD cohort. Red pluses indicate mean residual age at onset for every sum of STR repeat lengths. Grey and black dashed lines are plotted using coefficients of the linear regression analysis and regression with selection.

The sum of QTR lengths was found to predict residual age at onset significantly better than the difference in QTR lengths, the minimum or maximum QTR length, or the number of copies of the 3-repeat allele (Additional file 1: Table S4, Methods). QTR lengths are thus likely to influence age at onset in an additive manner.

The relationship of the association between residual age at onset and the sum of QTR repeat lengths with those of neighbouring SNVs is shown in Fig. 4 for the 468 individuals with both SNV and sequencing data. In these individuals, the significance of the association between residual age at onset and sum of repeat lengths (p = 1.2×10^−7^) was greater than that observed with the most significant SNV, rs79727797 (p = 3.6×10^−5^). To determine whether the sum of the QTR lengths or rs79727797 was driving the association with age at onset, we performed a conditional analysis in the 468 individuals with both SNV and sequencing data. When the association of rs79727797 with residual age at onset was conditioned on the sum of the QTR lengths, the p-value in our sequenced cohort dropped from p = 3.6×10^−5^ to p = 0.83. However, conditioning the association of age at onset with the sum of QTR lengths on rs79727797 genotypes, it remained significant (p = 9.2×10^−4^), indicating that the hexanucleotide QTR, and not rs7977797, is likely to be driving the signal in our data (Fig. 4). Manhattan plots of SNV associations with residual age at onset for the 468 individuals with SNV data, conditioning on the sum of QTR lengths and rs7977797 in turn, are shown in Additional file 1: Fig. S3.

**Fig. 4.**
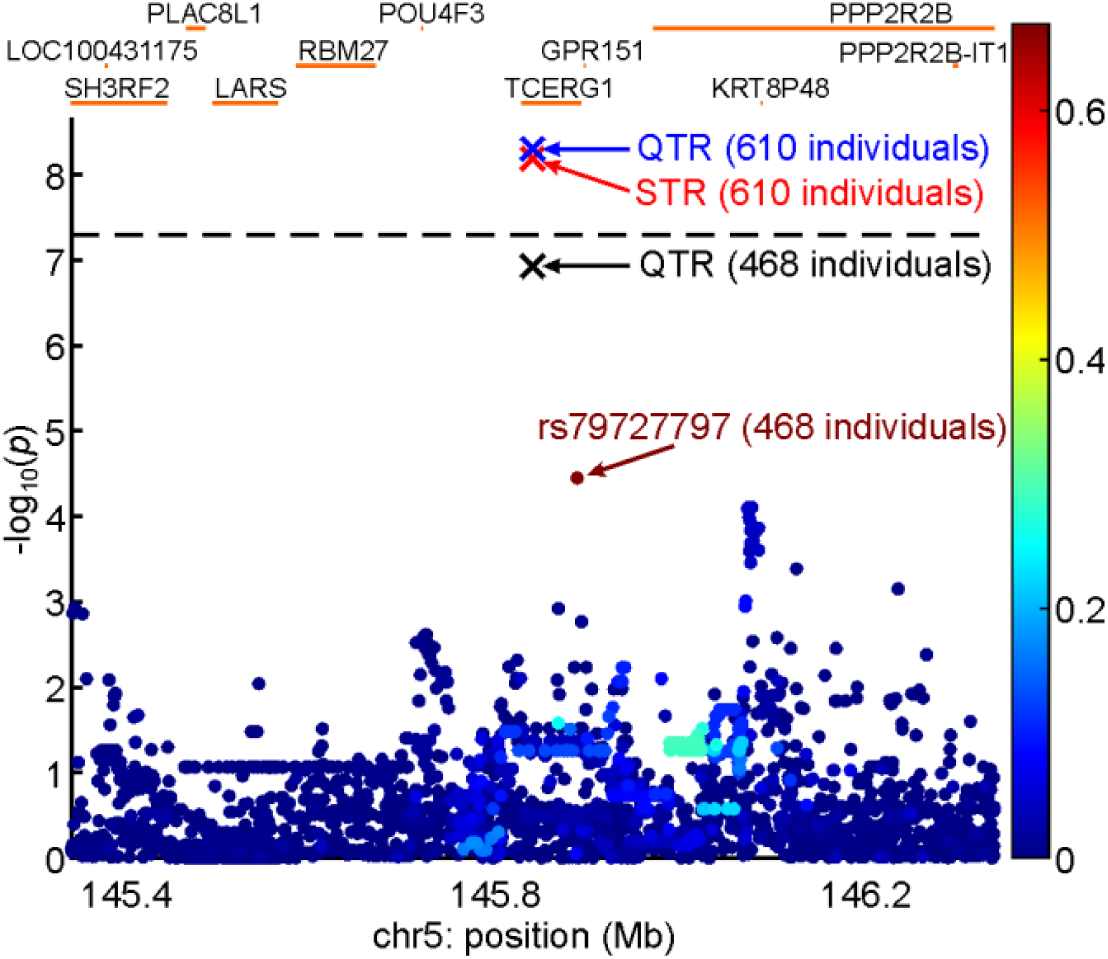
Locus zoom plot showing the relationship of rs79727797 association with residual age at onset to that of the sum of two short tandem repeat (STR) lengths (black cross) in 468 subjects with both single nucleotide variant (SNV) and sequencing data. The associations of age at onset with the sum of STR (red cross) and QTR (blue cross) repeat lengths in all 610 subjects are also shown. The bar on the right of the plot indicates the strength of linkage disequilibrium (r2) between each SNV and the tandem repeat. The p-value threshold for genome-wide significance (5×10^−8^) is shown with a black dashed line.

### Gene expression analyses

*TCERG1* has significant cis-expression quantitative trait loci (eQTLs), which can be used in conjunction with GWAS data to predict gene expression [18] in several tissues: GTeX [19] whole blood, PsychEncode [20] cortex, and eQTLGen whole blood [21]. rs79727797 is significantly associated only with expression of the nearby gene *PPP2R2B* (expansions in which cause SCA12) in eQTLGen (p=1.13×10^−16^), with the A allele that is associated with later onset being associated with increased expression of *PPP2R2B*. However, there are several SNVs more significantly associated with *PPP2R2B* expression in eQTLGen, and these have only modest significance in the GeM-GWAS (p-values of ∼0.07, see Additional file 2: Table S5). Likewise, the most significant eQTL SNVs for *TCERG1* in eQTLGen are not associated with HD age at onset in GeM (Additional file 3: Table S6). Notably, rs79727797 is not significantly associated with *TCERG1* expression (p=0.45). This indicates that gene expression (at least in whole blood) is unlikely to be the mechanism through which *TCERG1* influences age at onset in HD. This is corroborated by summary Mendelian Randomisation analyses using the eQTLGen expression data, which were non-significant (p=0.974 for *TCERG1*, p=0.07 for *PPP2R2B*). Co-localization analyses further showed that the eQTL and GWAS signals were different for both genes (colocalization probability=0). The lack of overlap between GeM GWAS association and eQTLGen and eQTL for *TCERG1* and *PPR2R2B* can be seen graphically in Additional file 1: Figs. S3 and S4.

We used FUSION [22] to perform TWAS analyses of the GeM dataset using the PsychENCODE [20] cortex expression data. There was a significant negative association between *TCERG1* expression and age at onset (Z=-2.71, p=0.00671): increased *TCERG1* expression is associated with earlier HD onset. Although the plot of eQTL and GWAS association (Additional file 1: Fig. S7) shows some overlap in signal, as does the table of significant eQTLs (Additional file 4: Table S7), a co-localization analysis does not show evidence that the eQTL and GWAS signals share the same causal variant (colocalization probability=0.0745). However, this analysis is inconclusive due to the relatively weak eQTL and GWAS signals (note that rs79727797 is not included in the analysis since the PsychENCODE sample is too small to demonstrate association with expression). No TWAS analyses were possible for *PPP2R2B*, since an insufficient proportion of variation in expression is attributable to SNVs. However, the plot of PsychENCODE eQTL and GWAS association (Additional file 1: Fig. S7) and table of significant eQTLs (Additional file 5: Table S8) show little overlap, which is supported by a colocalization analysis (colocalization probability=0.0376).

## Discussion

*TCERG1* is the only previously detected candidate gene for modifying HD age at onset to be confirmed by genome-wide association [8]. Our conditional analysis is consistent with the hexanucleotide tandem repeat in exon 4 explaining the signal attributed to the GWAS-significant SNV rs79727797 (which tags the three-repeat allele A2). The strength of the effect is directly proportional to the repeat length of the *TCERG1* QTR, with shorter repeats associated with later onset and longer repeats with earlier onset of HD. The previous finding that a slightly earlier than expected age at onset was detected in individuals whose longest allele is one and half hexanucleotide repeats longer than the reference [10] is consistent with our results (the participants with the genotype (38,40) in Fig. 2A most likely correspond to the inaccurately sized genotype (38,39.5) in [10]). The effect of the number of hexanucleotide repeats appears to be additive with each additional repeat giving one year earlier onset of HD: sum of repeats is significantly better associated with age at onset than either individual repeat allele or the difference between them (Additional file1: Table S4). The previous study [10] did not find that fitting the combined length of the two alleles improved the significance of the association with age at onset but did not formally compare the various models for allele length. That we were able to show a significant difference is likely due both to a larger sample, in which power was further increased by sampling individuals with extreme ages at onset, and to testing repeat lengths directly rather than allele lengths. Given the GWAS significant signal at this locus in an unselected HD population [8] we expect that this finding will replicate in unselected HD patients. Replication through sequencing the hexamer repeat in a larger unselected cohort is needed to assess the true effect size and the relationship of the modifier effect to repeat length.

*TCERG1* has known functions in transcriptional elongation and splicing [12,13]. It is in the top 5% of genes most intolerant of missense mutations, suggesting an essential role in cell biology [14]. How the *TCERG1* hexanucleotide repeat length modifies HD onset is unknown. Possibilities include *cis* or *trans* modulation of *TCERG1* or other gene expression, modulation of RNA splicing or transcription-splicing coupling, and effects on somatic expansion of the CAG repeat in *HTT*. Effects could be mediated by the tandem repeat in DNA or RNA, or by the translated (QA)_n_ tract in protein. The QTR has a slightly stronger association signal than the central STR, which may reflect an association with the length of the QA repeat in the protein rather than the CAGGCC hexamer in the DNA but more work is required to substantiate this observation. In DNA, repeat loci can modulate gene expression in *cis* [23,24], while transcribed repeats in RNA, especially tri- and hexamer repeats, can alter splicing, associate with R-loops and alter RNA stability or binding [25]. The hexamer repeat in *TCERG1* could act via altering expression of *TCERG1* or the nearby gene *PPP2R2B*. In our analysis evidence for the involvement of *TCERG1/PPP2R2B* expression in modification of HD age at onset is unclear. It was not possible to test the association of the *TCERG1* repeat with expression directly and the tagging SNV (rs79727797) is relatively rare (minor allele frequency = 2.4%), so requires a very large expression sample to show any association. Only eQTLGen (whole blood) is sufficiently large (n=31,684), and in this sample rs79727797 is significantly associated with *PPP2R2B* rather than *TCERG1* expression. However, the summary Mendelian Randomisation analyses are not significant for either gene, suggesting that neither *TCERG1* nor *PPP2R2B* expression is causally involved in modifying age at onset in HD, at least in blood. A significant TWAS association was observed in the PsychENCODE cortex expression data between reduced *TCERG1* expression and later age at onset, although there was little evidence that the eQTL and GWAS signals were co-localized. However, rs79727797 was not part of the TWAS predictor, due to the insufficient size of the PsychENCODE eQTL dataset. This weakened the GWAS signal, and thus reduced the power of the co-localization analysis. Furthermore, it was impossible to perform TWAS or co-localization analyses in caudate or striatum due to the lack of suitable eQTL datasets (the GTEx caudate sample is too small to show eQTL association with *TCERG1*). Hodges et al. [26] did not observe significant differential expression of *TCERG1* between HD patients and controls in caudate, although this study assessed expression via microarrays rather than more modern techniques. Langfelder et al. [27] observed significantly increased *TCERG1* expression in the striata of Q111, Q140 and Q175 mice relative to wild type. However, this has been suggested to be a compensatory homeostatic response to promote neuron survival [28], and such an effect would be difficult to model in a human eQTL sample. Therefore, it is possible that reduced *TCERG1* expression is associated with later onset of HD but corroborating evidence from other samples or direct experimentation is required for confirmation. Consistent with the observations of Langfelder et al.[27], immunostaining of post-mortem human brain showed increased nuclear *TCERG1* in HD caudate and cortex compared with normal controls, and increased staining with HD grade, suggesting that there may be a localisation effect of the repeat as suggested previously [15] and that excess nuclear *TCERG1* is deleterious in HD [10].

The hexanucleotide tandem repeat in *TCERG1* encodes an imperfect (QA)_n_ repeat in the protein and there are conflicting data on the role of this repeat in modulating normal *TCERG1* function. One reporter assay found the QA repeat to be dispensable for TCERG-mediated transcriptional repression [15], whereas a larger study in two cell lines found the QA repeat to be required for *TCERG1*-induced repression of the C/EBPα transcription factor [29]. A minimum of 17 QA repeats was required for this activity. When the QA repeat was deleted ΔQA-*TCERG1* colocalised with wild-type *TCERG1* and prevented its canonical relocalisation from nuclear speckles to pericentromeric regions, implicating a possible dominant negative mode of action. This is consistent with the QA repeat being required to retain the nuclear localisation of *TCERG1* [15], though not for its effect on transcription, although these overexpression experiments do not distinguish the effects of DNA, RNA and protein. A dominant negative mode of action would be inconsistent with the additive genetic effect we observe, although the effects we see relate only to differences of up to 5 units of the QA repeat in each *TCERG1* allele, rather than a complete deletion of the QA tract. Effects of this smaller modulation in the QA repeat are therefore likely to be more subtle. Taken together with the evidence that increased nuclear localisation of *TCERG1* is seen in HD mouse brain [27] it is plausible that the alteration in nuclear localisation conferred by the repeat could be responsible for the observed effect of *TCERG1* on age at onset. It remains possible that *TCERG1* expresses a novel function in cells with an expanded repeat unrelated to its normal function.

Many of the known genetic modifiers of age at onset of HD are proteins that act on DNA, particularly those involved in mismatch repair. These appear to operate by altering the levels of instability and expansion of the *HTT* CAG repeat, though there is also evidence for wider DNA repair deficits in HD [30,31]. It is possible that *TCERG1* modifies HD onset by acting directly or indirectly on the mechanisms regulating somatic expansion. Expansions of the inherited *HTT* CAG length are most marked in non-dividing neurons, suggesting that these events take place during transcription or DNA repair. *TCERG1* affects the processivity of RNA polymerase and splicing events during transcription, especially co-transcription [12,13]. During co-transcription it appears to bind and dissociate from stalled spliceosome complexes transiently [13] and the QA repeat might modulate this transient binding as it does with the C/EBPα interaction [29]. *HTT* exon 1 contains an RNAPII pause site [32], associated with co-transcriptional splicing [33–35]. Pausing associated with co-transcriptional splicing of *HTT* could stabilise the DNA-RNA hybrid R-loops that occur during active transcription [36–38]. Stabilised R-loops would give opportunities for increased binding and processing by the DNA repair machinery, and promote somatic expansion of the CAG repeat in *HTT* exon 1. Pausing might also promote aberrant splicing of *HTT* exon 1 which is regulated by RNAPII transcription speed [39]. This would likely generate a vicious cycle as lengthening repeats lead to increased RNAPII pausing followed by further dysregulation of exon 1 splicing and production of toxic exon 1 HTT species [40]. Stabilised R-loops are also associated with increased levels of DNA breaks in CAG/CTG repeats cleaved by MuLγ, encoded by *MLH1*/*MLH3*, both associated with modulating the length of CAG and other expansions [41– 43]: *MLH1* is associated with altered age at onset of HD [44]. Of note, knockdown of *TCERG1* in HEK293T cells leads to dysregulation of over 400 genes, including downregulation of *MLH1* [12].

The role of *TCERG1* in transcription could signal its involvement in the widespread transcriptional dysregulation that is seen in HD [12,26,45]. *TCERG1* is involved in the assembly of small nuclear ribonucleoproteins in mRNA processing [46]. It also interacts with huntingtin [10]. In yeast, proteins containing a (QA)_15_ tract can bind to a fragment of mutant huntingtin containing 103 glutamines to suppress its toxicity [47]. In amyotrophic lateral sclerosis and some cases of frontotemporal dementia, *TCERG1* increases the levels of TDP-43, the major constituent of the pathological hallmark inclusions in mammalian cells [48]. Notably, TDP-43 is observed alongside mHTT in extranuclear pathogenic inclusions in HD [49]. The genetic association of the CAGGCC/QA repeat in *TCERG1* with age at onset of HD is robust, with a hint that it might operate at level of the protein rather than DNA. More work is needed to clarify the mechanism by which it alters onset in HD and whether this is related to previously reported pathophysiologies or a new pathway. It provides a further potential treatment target in this incurable disease.

## Conclusions

We have identified a variable hexanucleotide QTR in *TCERG1* as a modifier of HD onset, with one year reduction in age at onset of HD for each additional hexamer repeat. Elucidation of the mechanism of its modifier effect will inform research into pathogenesis in HD and, potentially, other repeat expansion disorders, and could identify new therapeutic targets.

## Materials and Methods

### Subject details

We analysed genetic and phenotypic data of 506 patients with HD from the EHDN REGISTRY study (http://www.ehdn.org; [16]; initially we had 507 individuals, but then we excluded one individual with unreliably called *TCERG1* QTR due to low sequencing depth coverage), and 104 individuals from the Predict study [50]. Ethical approval for Registry was obtained in each participating country. Investigation of deidentified Predict-HD subjects was approved by the Institutional Review Board of Partners HealthCare (now Mass General Brigham). Participants from both studies gave written informed consent. Experiments were conducted in accordance with the declaration of Helsinki and ethical approval was Cardiff University School of Medicine SMREC 19/55.

DNA of the 506 REGISTRY HD individuals was provided by BioRep Inc. (Milan, Italy) from low-passage lymphoblastoid cells. For most of our HD patients (496 individuals), we measured the length of uninterrupted *HTT* exon 1 CAG repeat using an Illumina MiSeq platform [51]. For the remaining 10 individuals, we used BioRep CAG lengths determined using Registry protocols (https://www.enroll-hd.org/enrollhd_documents/2016-10-R1/registry-protocol-3.0.pdf). For individuals from the Predict study, DNA was obtained from blood DNA and we used the CAG length recorded in the study. SNV genotype data were available for 468 of the REGISTRY individuals, as part of the GeM GWAS[8].

Age at onset was assessed as described in [11]. For REGISTRY age at motor onset data, where onset was classified as motor or oculomotor by the rating clinician, the clinician’s estimate of onset was used for onset estimation. For all other onset types, we used the clinical characteristics questionnaire for motor symptoms. Predict age at motor onset was as recorded in the study, determined using the age where the diagnostic confidence level = 4. The selection of the REGISTRY and Predict samples are described in detail in [11]. Briefly, the REGISTRY samples were selected for having extreme early or late onset compared to that predicted by their CAG length. The Predict-HD samples were selected based on extreme predicted early or late onset. These originally constituted 232 individuals, of whom we analysed on those 104 who had a known age at motor onset.

### Calling tandem hexamer from whole exome sequencing (WES) data

For the Registry-HD cohort (N=506), sequencing was performed at Cardiff University [1]. Whole-exome libraries were generated using TruSeq® rapid exome library kits (Illumina, 20020617) according to Illumina protocols (https://emea.support.illumina.com/downloads/truseq-rapid-exome-library-prep-reference-guide-1000000000751.html). Libraries were sequenced on a HiSeq 4000 using 75bp paired-end reads. For the Predict-HD participants, an in-solution DNA probe based hybrid selection method was used to generate Illumina exome sequencing libraries. A HiSeq 2500 was used to generate 76bp paired end reads. De-multiplexed reads for both sets of exomes were aligned using BWA v0.7.5a [52], generating variant-ready binary alignment (BAM) files which were used for STR/QTR calling. Individuals with more than one sequencing run were merged into a single BAM file. The human genome assembly hg19 was used for sequence alignment. The genotyping was performed using universal variant caller (UVC) software openly available at https://github.com/LobanovSV/UVC.git. This software allows to call SNPs, INDELs, STRs, QTRs and any their combination in several steps: (i) align single reads to the reference genome using unique matching algorithm; (ii) remove reads with bad alignment score; (iii) find all possible combinations of insertions and deletions appearing from single read alignment; (iv) re-align reads to all of those combinations and choose the best one (i.e. having the smallest mismatch error); (v) correct alignment of single reads using their pairs as well as other reads; split reads into two (if possible) groups of similar size but with different allele sequence; correct read alignment using alignments of other samples. To align single reads to the reference genome, we construct the match matrix (Additional file 1: Fig. S8) and select the path minimising the mismatch error 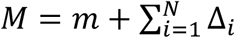 exp(5 −*l*_*i*_). Here, *m* is sum of mismatch nucleotides, *N*is number of gaps, Δ_*i*_ is the gap height (the distance between two match pieces adjoining the *i*-th gap), and *l*_*i*_ is minimal length of two match pieces. The mismatch error of this form takes into account the highly mutative nature of STRs/QTRs and allows to unbiasedly align reads with any combination of SNPs, INDELs, STRs and/or QTRs. For instance, the naive straight red line in Additional file 1: Fig. S8 has the mismatch error *M* = 2 (*m* =2, *N* = 0), whereas the correct blue path with 3 hexamer deletion has much smaller mismatch error *M* = 6.2 · 10^−13^ (*m* = 0, *N* = 1, Δ_*i*_= 18, *l*_*i*_ = 36). If the right adjoining piece is located above the left one (for example, the break of the blue line in Additional file 1: Fig. S8), the gap is attributed as a deletion. Conversely, the discontinuity is attributed as an insertion. Finally, we create an alignment track with rows containing sequences of mapped paired reads. To simplify genotyping, we expand the sequence of the reference genome by inserting asterisks to the loci at which the reads have insertions. Conversely, we substitute nucleotide deletions in the reads by asterisks. This manoeuvre permits insertions and deletions to be treated as substitutions. After that we consider loci where some sequence reads have nucleotides different from the reference. We utilise these loci to retrieve the allele sequences by separating the reads into two groups in such a way that all reads in a single group have the same nucleotides at these loci.

### Sanger sequencing to confirm QTR sequences

To validate our tandem hexamer calls from WES data, we performed Sanger sequencing of four samples: two homozygous for the reference QTR allele (A1/A1 genotype), one heterozygous for a shorter QTR allele (A1/A2 genotype), and one heterozygous for a longer QTR allele (A1/A4 genotype). The QTR locus in *TCERG1* was amplified by PCR using forward (5’-AACTGACACCTATGCTTG-3’) and reverse (5’-GTTGAAGTGGATACTGCA-3’) primers as described in the reference [10]. Amplicons were Sanger sequenced (LGC, Germany) in both directions using forward (5’-AACTGACACCTATGCTTGCAG-3’) and reverse (5’-GAAGTGGATACTGCAGGTGC-3’) primers, and sequences compared to their respective calls from short-read exome sequencing data. Sequences from Sanger and exome sequencing matched in each of the four cases.

### Measuring *TCERG1* QTR lengths using capillary electrophoresis

To confirm *TCERG1* QTR lengths derived from exome-sequencing data, the QTR locus in *TCERG1* was amplified by PCR using a fluorescently-labelled forward (5’-FAM-AACTGACACCTATGCTTG-3’) and unlabelled reverse (5’-GTTGAAGTGGATACTGCA-3’) primer before sizing by capillary electrophoresis (ABI 3730 genetic analyzer) and Genescan against a LIZ600 ladder of size standards (Thermofisher). In total we tested QTR length calls for 101 individuals from the Registry-HD sample: the 73 who had at least one non-reference QTR length allele (A2-A8) and 28 who were called as homozygous for the reference (A1) allele. The reference allele A1 was predicted to produce a PCR fragment of 307 bp. In all samples this allele was consistently sized at 299 bp by capillary electrophoresis. We attributed this to the repetitive nature of the sequence and the specific analyzer used. In all 101 individuals tested, allelic QTR lengths relative to the reference A1 allele QTR length exactly matched those called using exome-sequencing data.

### Calculation of age at onset residuals

Expected ages of onset were calculated from patient CAG length data (measured as described above) using the Langbehn model [53]. Residual ages at motor onset were then calculated taking the difference between the expected onset from the recorded clinical age at motor onset, as performed elsewhere [8].

### Association of age at onset with STR/QTR repeats

Linear regression was performed of the age at onset residual on the repeat statistic (sum, diff, max, min, #3 rep). Since the sample was selected to have extreme values (positive and negative) of this residual, linear regression is likely to overestimate the effect of the repeat on onset in the general HD patient population. Therefore, regression with selection (see Additional file 1: Supplementary Methods) was used to estimate the true effect size. A dichotomous phenotype was derived by selecting individuals with extreme late (positive residual greater than a pre-defined criterion) or early (negative residual less than a pre-defined criterion) onset.

Association of the dichotomous phenotype with repeat statistic was tested via logistic regression.To formally test which repeat statistics best predict age at onset, we proceeded as follows: For each pair of statistics A and B, a linear regression of residual age-at-onset on statistic A was performed as a baseline. Then statistic B was added to the regression and the significance of the improvement in fit assessed using ANOVA. Statistics were defined as “best fitting” if the addition of no other statistic gave a significant improvement in fit.

### Analyses to test for correlation between genetically predicted expression and age at onset

FUSION[22] was used to perform TWAS analyses on the PsychENCODE data using pre-computed predictors downloaded from http://resource.psychencode.org/. Summary Mendelian Randomisation was used to perform TWAS analyses on eQTLGen blood expression using cis-eQTL data downloaded from https://www.eqtlgen.org/cis-eqtls.html. Co-localisation analyses to test if eQTL and age at onset signal share the same causal SNV were performed using COLOC [54].

## Abbreviations

STR: Short tandem repeat
QTR: Quasi-tandem repeat
SNV: Single nucleotide variant
GWAS: Genome-wide association study
LD: Linkage disequilibrium
MAF: Minor allele frequency
eQTL: Expression quantitative trait locus
HD: Huntington’s disease

## Acknowledgements

We thank all the patients who contributed data to this research. We thank the Institute of Medical Genetics for use of the ABI 3730 Genescan allele sizing service. S.L., B.Mc. were supported by CHDI, B.Mc. and S.L. by ARUK, B.Mc. by a PhD studentship from Cardiff University School of Medicine. J.F.G. and M.E.M. received support from NIH grant NS091161 and from the CHDI Foundation, Inc. J-M.L. received support from grant R01NS105709. A.E.R. received support from MRC, Wellcome Trust, Campaign for Alzheimer’s Research in Europe, Horizon 2020, JPND and Health and Care Research Wales. L.J., N.M.W. and P.H. were supported by an MRC Centre grant (MR/L010305/1). L.J., P.H., T.H.M. and N.D.A. were supported by CHDI. T.H.M. was supported by a Welsh Clinical Academic Track Fellowship, an MRC Clinical Training Fellowship (MR/P001629/1) and a Patrick Berthoud Charitable Trust Fellowship through the Association of British Neurologists. B.Mc., T.H.M and L.J. were supported by Brain Research Trust (No.201617-06). The two of the authors of this publication are a members of the European Reference Network for Rare Neurological Diseases - Project ID No 739510. This work was performed using the computational facilities of the Advanced Research Computing @ Cardiff (ARCCA) Division, Cardiff University. The Registry study is supported by the European Huntington’s Disease Network (EHDN), funded by CHDI Foundation, Inc.. The funding source had no role in study design; in the collection, analysis and interpretation of data; in the writing of the report; and in the decision to submit the paper for publication. We thank Vincent Dion for commenting on the manuscript.

## Authors’ contributions

Conceptualisation: T.H.M., L.J.; Methodology: S.L., B.Mc.; Software: S.L.; Validation: M.Mc.K., T.H.M.; Formal Analysis: S.L., P.H.; Investigation: S.L., B.Mc., M.Mc.K.; Resources: B.Mc., G.B.L., A.E.R., M.O., EHDN Registry, J.S.P., Predict-HD, J.-M.L., M.E.M., J.F.G.; Data curation: S.L.; Writing – original draft: S.L., P.H., L.J; Writing – review and editing; J.-M.L., M.E.M., J.F.G., J.D.L., M.R., T.H.M.; Visualisation: S.L.; Supervision: T.H.M., P.H., L.J., N.M.W.; Project administration: T.H.M., L.J.; Funding acquisition: L.J., P.H, T.H.M., M.R., N.M.W.

## Supplementary Information

**Additional file 1: Supplementary Information, Table S1**. Significance of the association between *TCERG1* exon 4 quasi-tandem repeat (QTR) and residual age at onset ℛ for the various ways of coding the repeat, **Table S2**. Significance of the association between the sum of *TCERG1* QTR lengths and residual age at onset in REGISTRY, Predict-HD and combined samples, **Table S3**. As Table S1, but for short tandem repeat (STR), **Table S4**. Significance (p-value) of improvement in fit to residual age at onset given by adding the “additional” QTR statistic to a model containing the “baseline” QTR statistic, **Fig. S1**. Transcription elongation regulator 1 isoform 3 [Homo sapiens], **Fig. S2**. As **Fig. 3**, but for short tandem repeat (STR), **Fig. S3**. Manhattan plots of residual age at onset association conditioning on rs79727797 and QTR, **Fig. S4**. Plots of *TCERG1* eQTL from eQTLGen and GeM GWAS, **Fig. S5**. Plots of *PPP2R2B* eQTL from eQTLGen and GeM GWAS, **Fig. S6**. Plots of *TCERG1* eQTL from PsychENCODE and GeM GWAS, **Fig. S7**. Plots of *PPP2R2B* eQTL from PsychENCODE and GeM GWAS. **Fig. S8**. Illustration of the match matrix. **Supplementary Methods**. Regression with selection.

**Additional file 2: Table S5**. List of significant eQTLGen eQTLs for *PPP2R2B* (see full description in **Additional file 1**).

**Additional file 3: Table S6**. List of significant eQTLGen eQTLs for *TCERG1* (see full description in **Additional file 1**).

**Additional file 4: Table S7**. List of significant psychENCODE eQTLs for *TCERG1* (see full description in **Additional file 1**).

**Additional file 5: Table S8**. List of significant psychENCODE eQTLs for *PPP2R2B* (see full description in **Additional file 1**).

## Availability of data and materials

The datasets supporting the conclusions of this article are included within the article and its additional files. The software performing regression with selection and STR/QTR calling are available from https://github.com/LobanovSV at the RegressionWithSelection and UVC repositories, respectively.

## Supplementary Information for

### Supplementary Tables

**Table S1.**
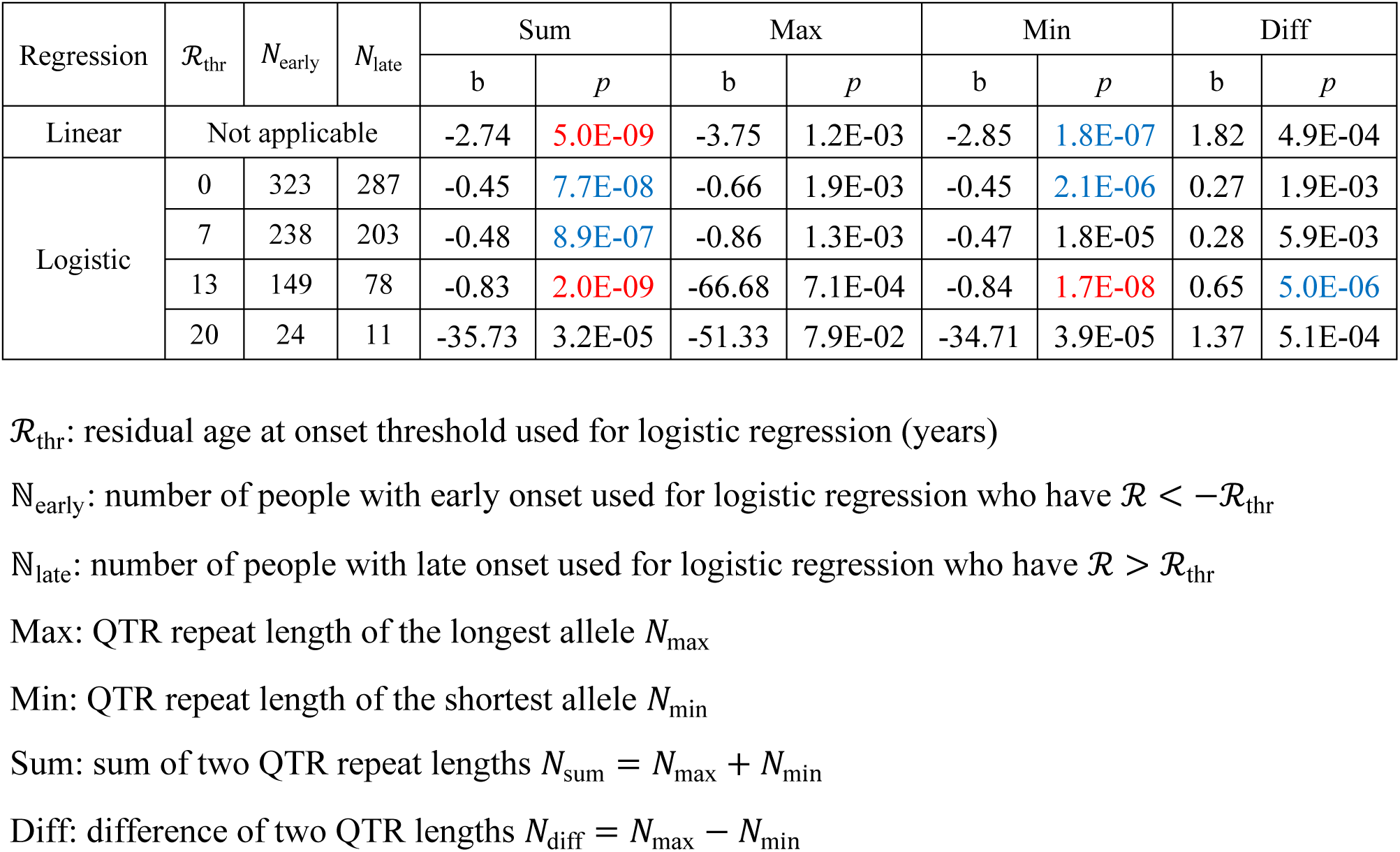
Significance of the association between TCERG1 exon 4 quasi-tandem repeat (QTR) and residual age at onset ℛ for the various ways of coding the repeat. Blue and red colours highlight numbers passing 10^−5^ and 5 · 10^−8^ *p*-value thresholds for significance, respectively.

**Table S2.** Significance of the association between the sum of TCERG1 QTR lengths and residual age at onset in REGISTRY, Predict-HD and combined samples. Two types of regression analysis of the REGISTRY cohort are presented: linear regression analysis and regression with selection (see Supplementary Methods section below).

| Cohort | Type of regression analysis of REGISTRY | b | SE | CI |  | *p* |
| --- | --- | --- | --- | --- | --- | --- |
| REGISTRY | Linear | -3.10 | 0.55 | -4.19 | -2.02 | 3.15E-08 |
|  | Selection | -0.98 | 0.19 | -1.36 | -0.62 | 2.30E-08 |
| Predict-HD | N/A | -1.26 | 0.56 | -2.37 | -0.14 | 0.027 |
| Combined | Linear | -2.74 | 0.46 | -3.65 | -1.84 | 5.02E-09 |
|  | Selection | -1.00 | 0.18 | -1.37 | -0.67 | 2.14E-09 |

**Table S3.** As **Table S1**, but for short tandem repeat (STR).

| Regression | $\mathcal{R}_{\text{thr}}$ | $N_{\text{early}}$ | $N_{\text{late}}$ | Sum | | Max | | Min | | Diff | |
| --- | --- | --- | --- | --- | --- | --- | --- | --- | --- | --- | --- |
| | | | | b | $p$ | b | $p$ | b | $p$ | b | $p$ |
| Linear | Not applicable |  |  | -2.75 | 6.5E-09 | -3.78 | 1.2E-03 | -2.85 | 2.4E-07 | 1.80 | 6.5E-04 |
| Logistic | 0 | 323 | 287 | -0.45 | 1.2E-07 | -0.64 | 2.5E-03 | -0.45 | 2.7E-06 | 0.27 | 2.0E-03 |
|  | 7 | 238 | 203 | -0.48 | 1.1E-06 | -0.86 | 1.3E-03 | -0.47 | 2.3E-05 | 0.28 | 7.4E-03 |
|  | 13 | 149 | 78 | -0.83 | 2.8E-09 | -66.68 | 7.1E-04 | -0.85 | 2.3E-08 | 0.65 | 7.3E-06 |
|  | 20 | 24 | 11 | -35.73 | 3.2E-05 | -51.33 | 7.9E-02 | -34.71 | 3.9E-05 | 1.37 | 5.1E-04 |

**Table S4.** Significance (p-value) of improvement in fit to residual age at onset given by adding the “additional” QTR statistic to a model containing the “baseline” QTR statistic. See **Table S1** for explanation of Sum, Max, Min, and Diff.

| Baseline model | Additional statistic |  |  |  |  |
| --- | --- | --- | --- | --- | --- |
|  | Sum | Max | Min | Diff | #3 repeats |
| Sum | X | 0.79 | 0.79 | 0.79 | 0.67 |
| Max | 1.07E-06 | X | 1.07E-06 | 1.07E-06 | 3.33E-05 |
| Min | 8.28E-03 | 8.28E-03 | X | 8.28E-03 | 0.71 |
| Diff | 2.61E-06 | 2.61E-06 | 2.61E-06 | X | 6.03E-03 |
| #3repeats | 1.29E-04 | 4.77E-03 | 5.49E-03 | 0.95 | X |

**Additional file 2: Table S5**. List of significant eQTLGen eQTLs for PPP2R2B with corresponding p-value for association with age at onset in the GeM GWAS. “P-value” = eQTLGen eQTL p-value, “Z-score” = eQTLGen test statistic. Positive Z means that the “assessed” allele is associated with higher expression. “AAO_effect” is the increase (or decrease, if negative) in age at onset (years) associated in the GeM GWAS with one copy of the “assessed” allele. P(GeM) is the p-value for association with age at onset in the GeM GWAS.

**Additional file 3: Table S6**. List of significant eQTLGen eQTLs for TCERG1 with corresponding p-value for association with age at onset in the GeM GWAS. “P-value” = eQTLGen eQTL p-value, “Z-score” = eQTLGen test statistic. Positive Z means that the “assessed” allele is associated with higher expression. “AAO_effect” is the increase (or decrease, if negative) in age at onset (years) associated in the GeM GWAS with one copy of the “assessed” allele. P(GeM) is the p-value for association with age at onset in the GeM GWAS.

**Additional file 4: Table S7**. List of significant psychENCODE eQTLs for TCERG1 with corresponding p-value for association with age at onset in the GeM GWAS. “eQTL_pval” = psychENCODE eQTL p-value, “eQTL_effect” = change in expression associated with each copy of allele A1. “AAO_effect” is the increase (or decrease, if negative) in age at onset (years) associated in the GeM GWAS with one copy of the “assessed” allele. P(GeM) is the p-value for association with age at onset in the GeM GWAS.

**Additional file 5: Table S8**. List of significant psychENCODE eQTLs for PPP2R2B with corresponding p-value for association with age at onset in the GeM GWAS. “eQTL_pval” = psychENCODE eQTL p-value, “eQTL_effect” = change in expression associated with each copy of allele A1. “AAO_effect” is the increase (or decrease, if negative) in age at onset (years) associated in the GeM GWAS with one copy of the “assessed” allele. P(GeM) is the p-value for association with age at onset in the GeM GWAS.

**Additional file 6: Table S9**. List of phenotypes and genotypes for the individuals with Huntington’s disease. NCAG: *HTT* CAG repeat length of the expanded allele

AAO: age at motor onset RAAO: residual age at onset

rs79727797: number of minor alleles (A) of the rs79727797 SNP

dNmin_QTR: *TCERG1* QTR repeat length of the shortest allele called from whole exome sequencing (WES) data

dNmax_QTR: *TCERG1* QTR repeat length of the longest allele called from WES data dNmin_STR: *TCERG1* STR repeat length of the shortest allele called from WES data dNmax_STR: *TCERG1* STR repeat length of the longest allele called from WES data dNmin_GeneScan: *TCERG1* QTR repeat length of the shortest allele measured using GeneScan dNmax_GeneScan: *TCERG1* QTR repeat length of the longest allele measured using GeneScan

### Supplementary Figures

**Fig. S1.**
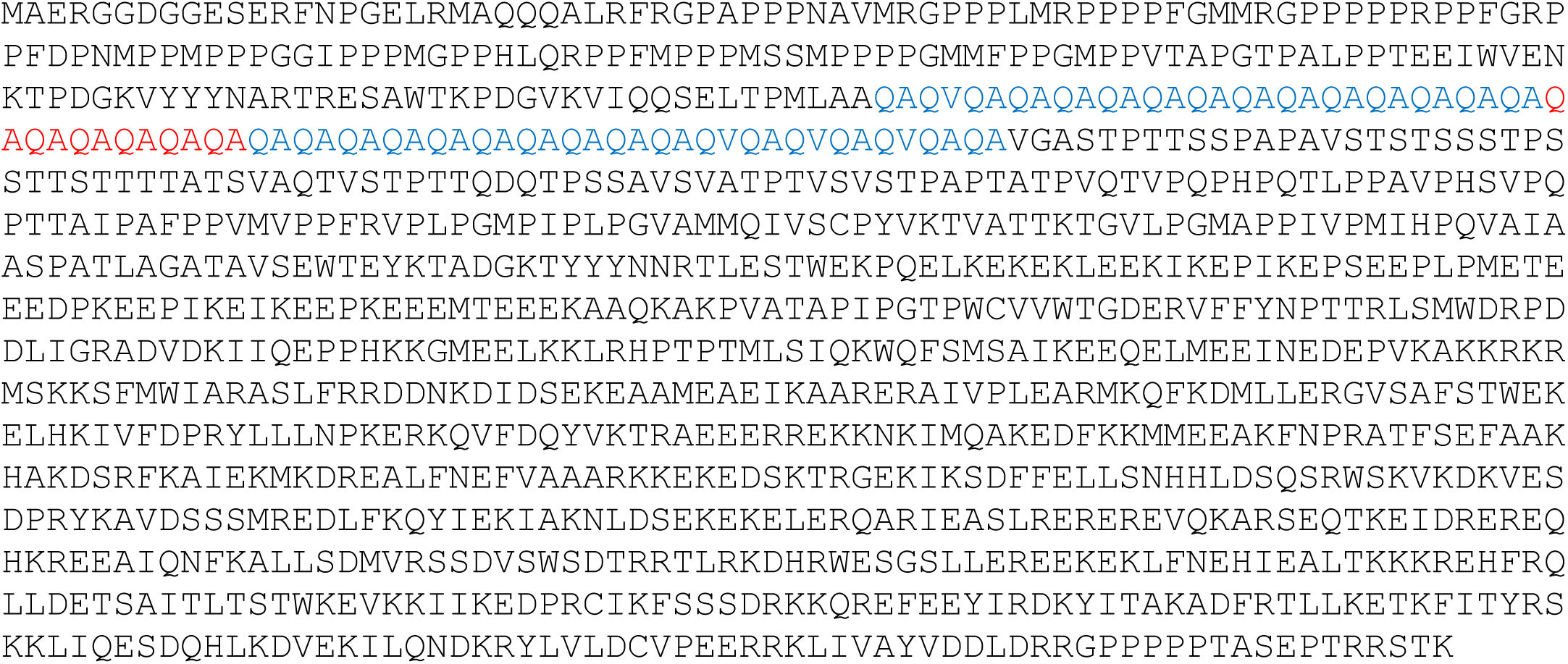
Transcription elongation regulator 1 isoform 3 [Homo sapiens]. NCBI Reference Sequence: NP_001369477.1

**Fig. S2.**
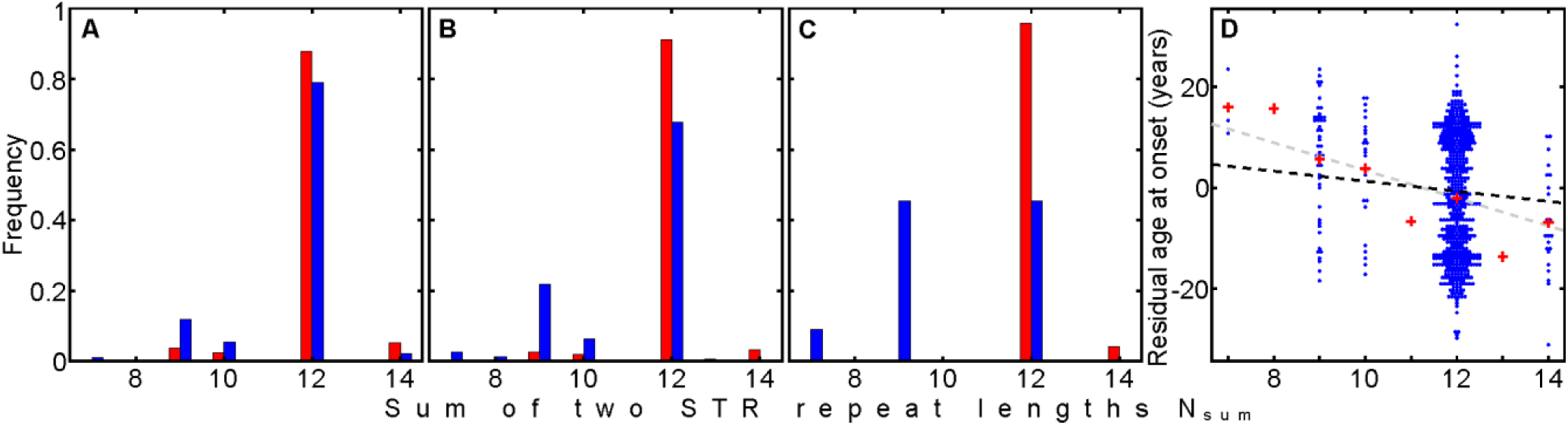
As **Fig. 3**, but for short tandem repeat (STR).

**Fig. S3.**
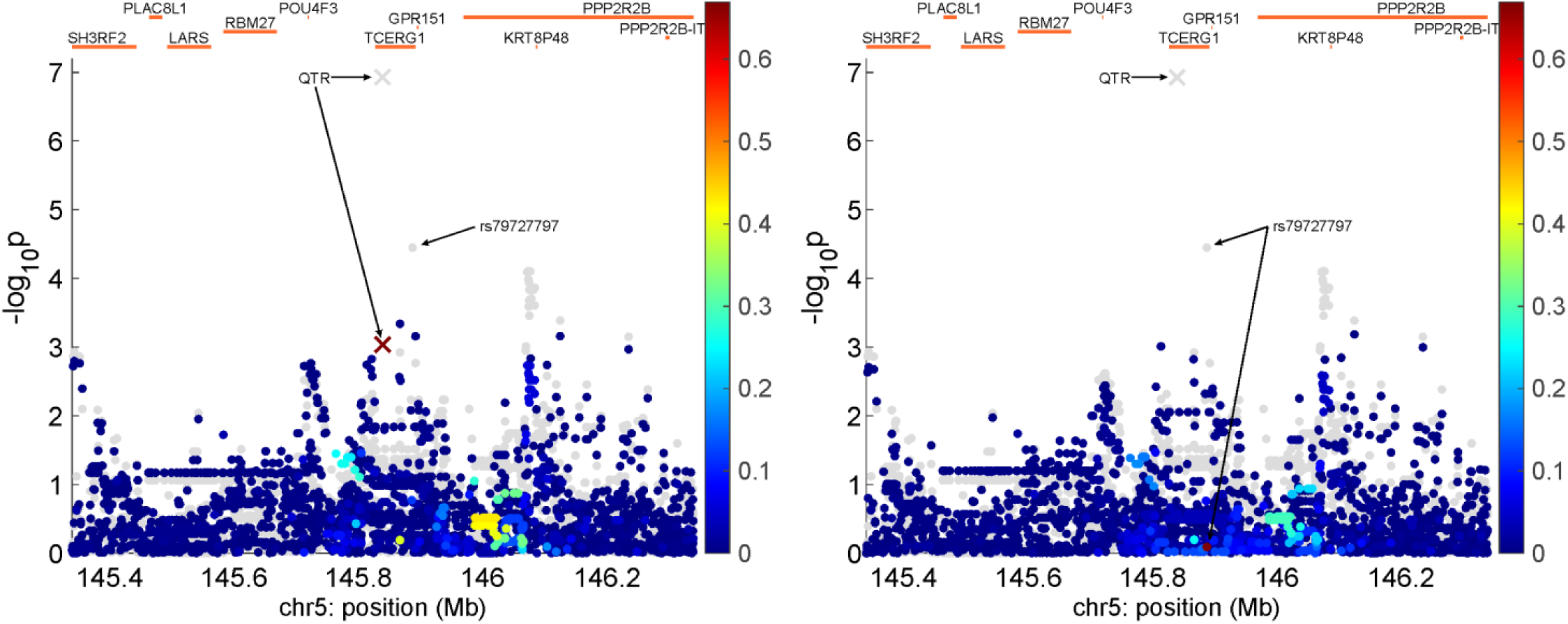
Manhattan plots of residual age at onset association conditioning on rs79727797 (left panel) and QTR (right panel) for 468 HD individuals with both sequencing and GWAS data. The bar on the right of the plots indicates the strength of linkage disequilibrium (r2) between each SNP/QTR and the variant being conditioned on. The grey dots mark p-values prior to conditioning. The variant being conditioned on necessarily disappears from the plot.

**Fig. S4.**
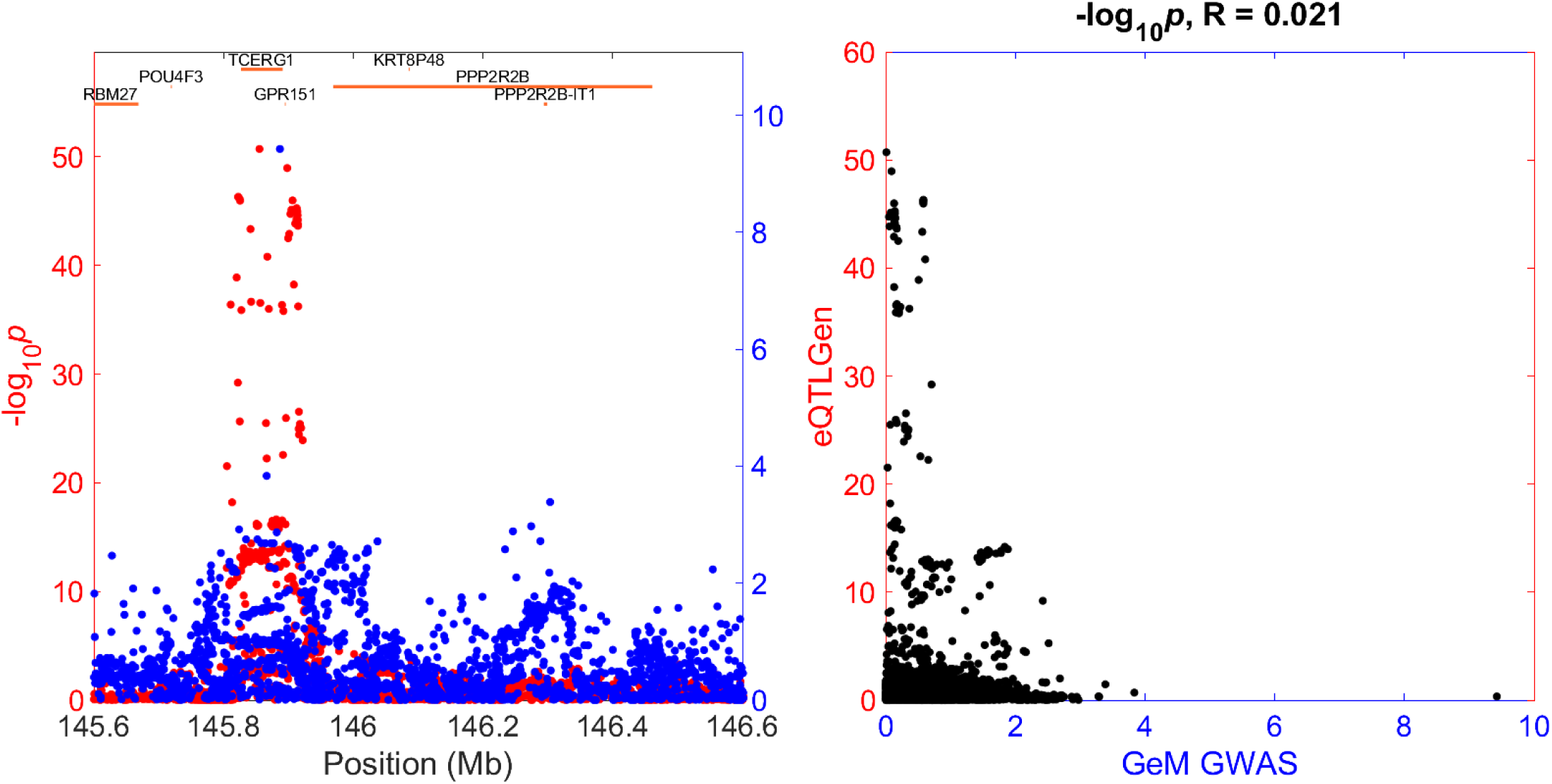
Plots of TCERG1 eQTL -log p-value from eQTLGen (red) and GeM GWAS -log p-value (blue) vs chromosome position (left panel) and each other (right panel)

**Fig. S5.**
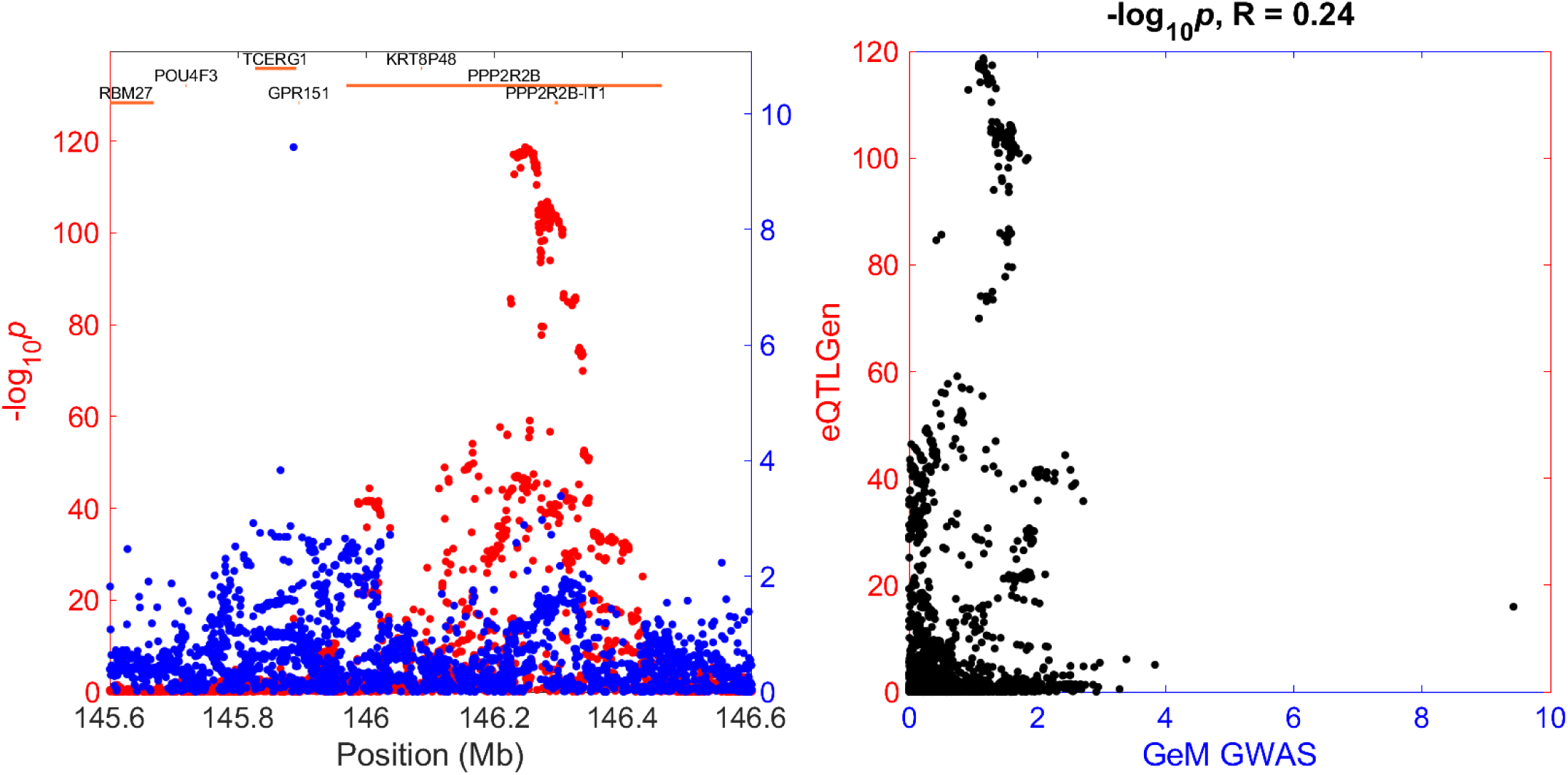
Plots of PPP2R2B eQTL -log p-value from eQTLGen (red) and GeM GWAS -log p-value (blue) vs chromosome position (left panel) and each other (right panel)

**Fig. S6.**
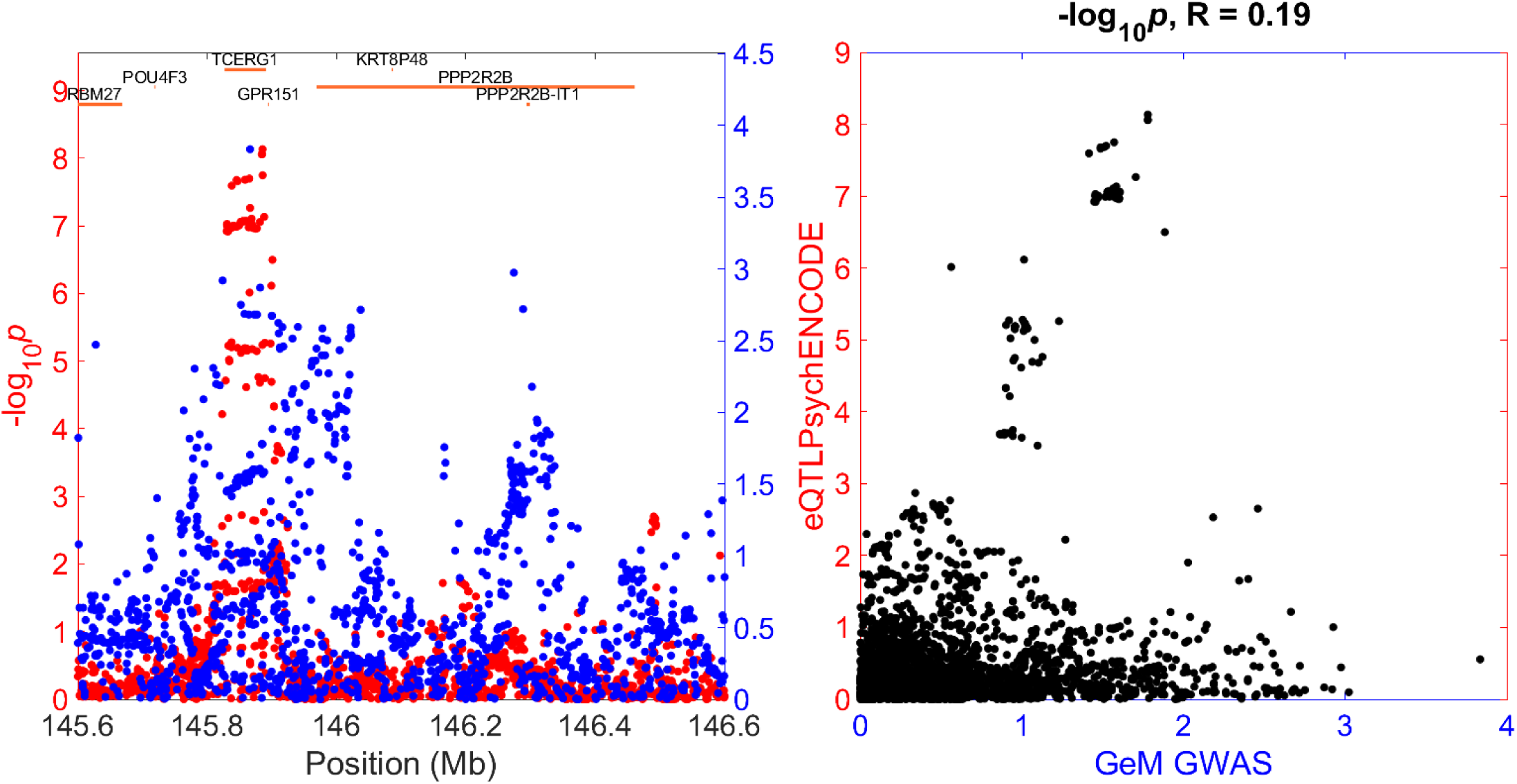
Plots of TCERG1 eQTL -log p-value from PsychENCODE (red) and GeM GWAS -log p-value (blue) vs chromosome position (left panel) and each other (right panel)

**Fig. S7.**
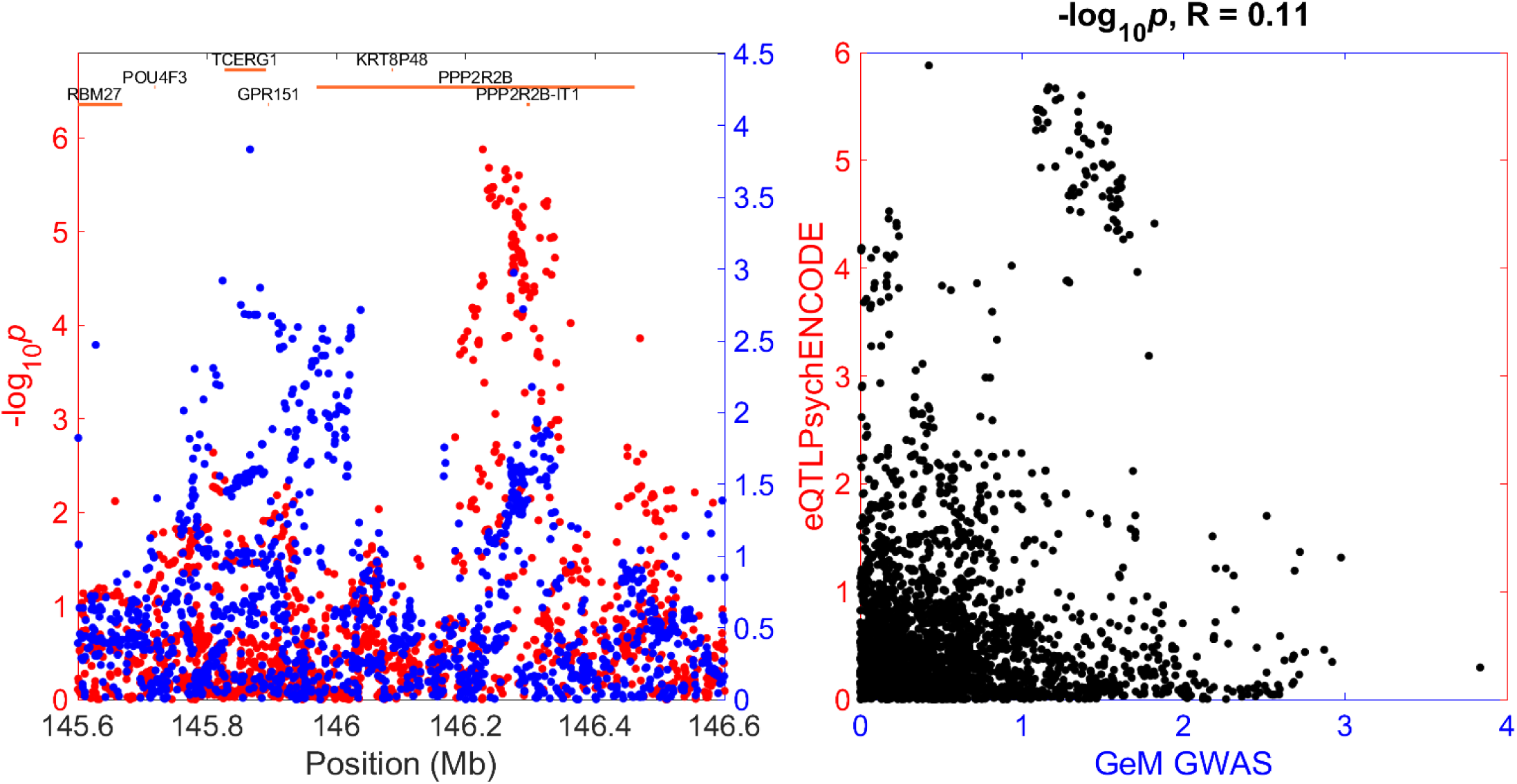
Plots of PPP2R2B eQTL -log p-value from PsychENCODE (red) and GeM GWAS -log p-value (blue) vs chromosome position (left panel) and each other (right panel)

**Fig. S8.**
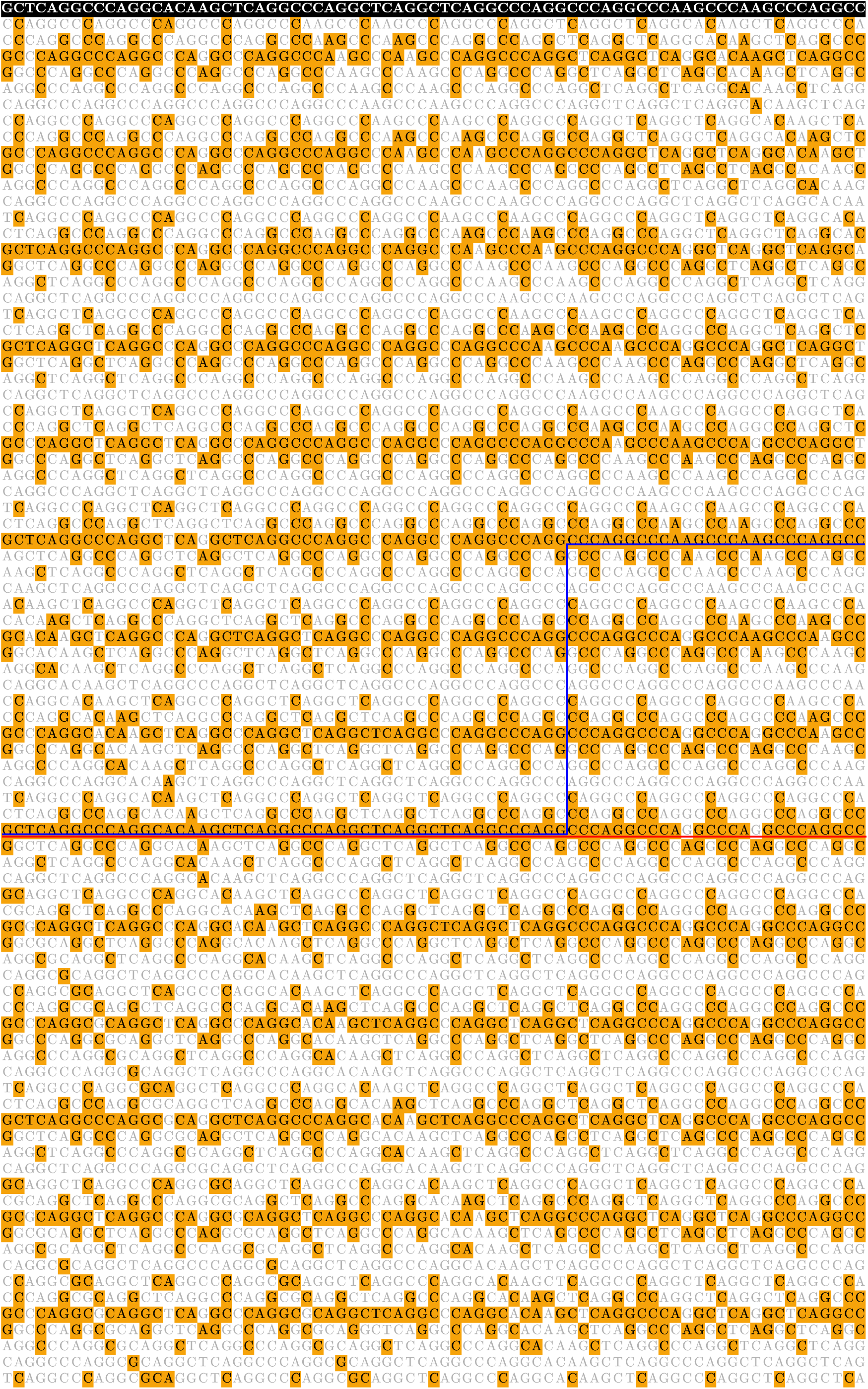
Illustration of the match matrix. The top line with white letters on a black ground specifies the nucleotide sequence of a read. Other lines represent nucleotide sequences of the reference genome shifted by one nucleotide with respect to the previous (upper) line. Matched nucleotides are highlighted with a yellow colour which creates the match matrix (yellow = True, white = False). Two read alignments are shown. The naive way to align a read with two mismatched nucleotides is shown as a red straight line. The path with one deletion, which takes into account the highly mutative nature of STRs/QTRs, is shown with a blue line.

## Supplementary Methods: Regression with selection

### Section S1: Initial selection

We performed whole-exome sequencing of a small sub-group of the EHDN REGISTRY study. To increase statistical power, we selected individuals with the largest absolute value of the residual age at onset *R*.

The probability density function of the residual ages at onset in the selected sample *p*(*R*) is that of a normal distribution with mean 0 and standard deviation σ

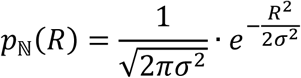

multiplied by a selection function

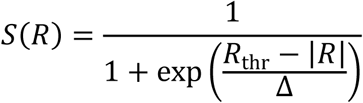

and normalised to have unitary integral

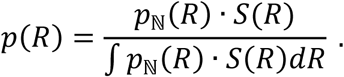

Here, σ is the standard deviation of the initial HD population (EHDN REGISTRY study), *R*_thr_ is the selection threshold, and Δis the selection width, which was infinitely small, Δ→ 0.

The expected probability density of the initial HD population *p*_ℕ_(*R*), selection function *S*(*R*), and expected probability density *p*(*R*) of the selected HD sub-group are shown in **Fig. S9**.

**Fig. S9.**
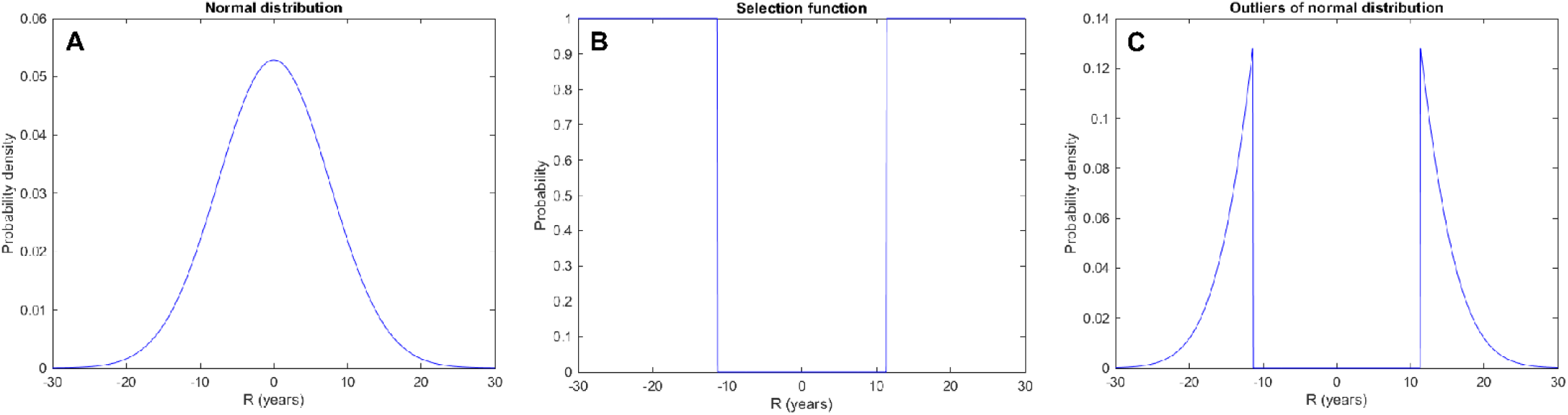
**a** Expected probability density of the initial HD population (normal distribution); **b** Selection function with infinitely small Δ; **c** Expected probability density of the small HD sub-group with largest absolute value of the residual age at onset |*R*|.

### Section S2: Correction of the age at onset residuals

To improve the accuracy of the correction of age at onset for CAG length, we additionally measured the length of the uninterrupted HTT exon 1 CAG repeat using an Illumina MiSeq platform for 496 individuals from our HD cohort and corrected the HD age at onset residuals. Some individuals who had age at onset residual above the threshold *R*_thr_ shifted to the region with |*R*| below the threshold *R*_thr_ after correction. Conversely, some individuals who would have corrected age at onset residual above the threshold *R*_thr_, were not selected because their uncorrected |*R*| were below the threshold *R*_thr_. The correction has therefore widened the selection function, corresponding to a non-zero selection width Δ.

The probability density function *p*(*R*) of our HD group with corrected residuals can be modelled in the same way as was described in **Section S1**, but with finite selection width Δ.

We estimated parameters of the selection function *S*(*R*) by minimising maximum absolute difference *D*

between the expected *F*_*e*_(|*R*|) and observed *F*_*o*_(|*R*|) cumulative probabilities

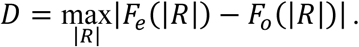

The observed cumulative probability *F*_*o*_(|*R*|) and expected one *F*_*e*_(|*R*|) with optimal parameters σ = 7.02, *R*_thr_ = 17.6, Δ= 3.30 are shown in **Fig. S10A**. The one-sample Kolmogorov–Smirnov *p*-value is 0.81. The selection and probability densities are shown in **Fig. S10B,C**.

**Fig. S10.**
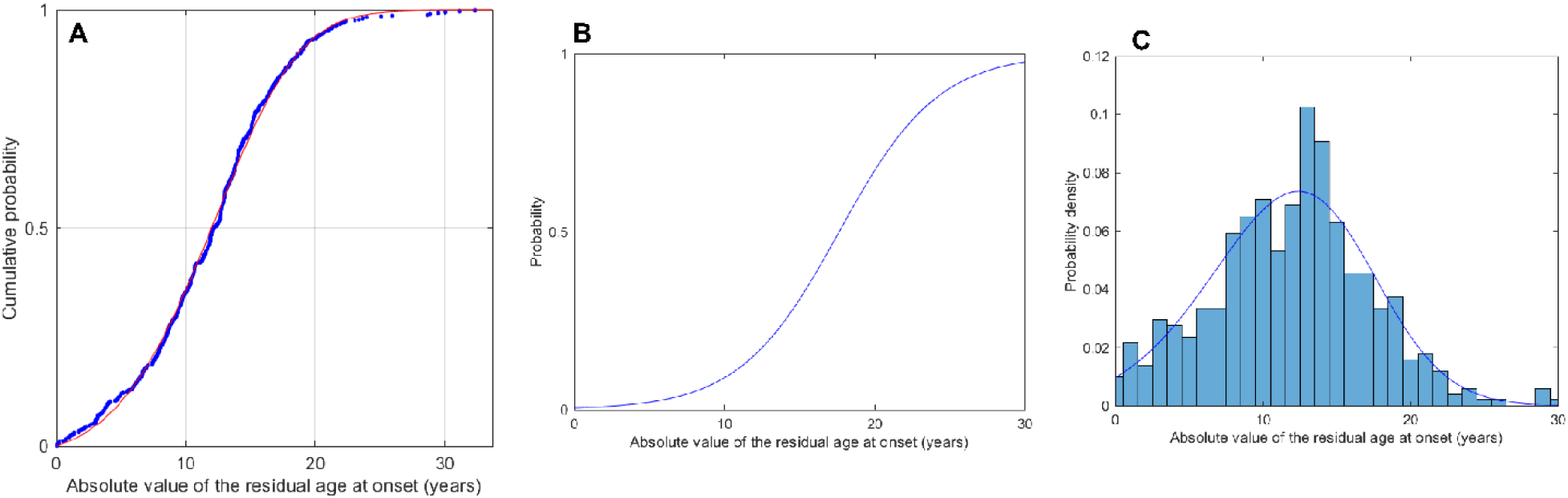
**a** The observed (blue dots) and expected (red line) cumulative probabilities; **b** The selection function with optimal parameters; **c** The observed (bars) and expected (line) probability densities.

### Section S3: Likelihood function

In the linear regression, the errors

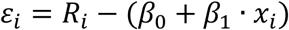

are normally distributed ε_*i*_∼ℕ(0, σ^2^) and are independent across individuals. The likelihood ℒ_LR_(σ, *β*_0_, *β*_1_|*R*_*i*_, *x*_*i*_) is

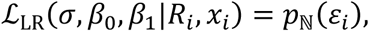

where *p*_ℕ_(ε_*i*_) is the probability density function of normal distribution (see **Section S1**). Here, σ, *β*_0_, and *β*_1_ are unknown standard deviation, intercept, and effect size, respectively; *R*_*i*_ and *x*_*i*_ are age at onset residual and sum of TCERG1 QTR lengths of a specific individual.

In the regression with selection, the distribution of the errors differs between individuals. The likelihood ℒ(σ, *β*_0_, *β*_1_|*R*_*i*_, *x*_*i*_) is

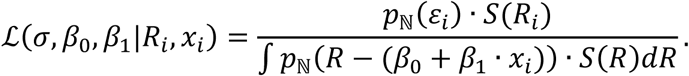

Note, the integral in the denominator depends on the individual’s sum of QTR lengths *x*_*i*_. Here, *S*(*R*) is the selection function (see **Section S1**).

We estimated the unknown parameters σ, *β*_0_, and *β*_1_ by maximising the likelihood function of all observations

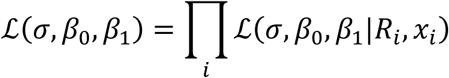

and used the likelihood-ratio test (comparing the likelihood maximised over σ, *β*_0_, *β*_1_ to that maximised over σ and *β*_0_, holding *β*_1_=0) to obtain the significance of the association.

### Section S4: Software

The software performing minimisation of the maximum absolute difference between the expected and observed cumulative probabilities (**Section S2**) and maximisation of the likelihood function (**Section S3**) is freely available from https://github.com/LobanovSV/RegressionWithSelection.git.

